# Preventing *S. aureus* biofilm formation on titanium surfaces by the release of antimicrobial β-peptides from polyelectrolyte multilayers

**DOI:** 10.1101/431205

**Authors:** Angélica de L. Rodríguez López, Myung-Ryul Lee, Riley Whitehead, David M. Lynn, Sean P. Palecek

## Abstract

*Staphylococcus aureus* infections represent the major cause of titanium based-orthopaedic implant failure. Current treatments for *S. aureus* infections involve the systemic delivery of antibiotics and additional surgeries, increasing health-care costs and affecting patient’s quality of life. As a step toward the development of new strategies that can prevent these infections, we build upon previous work demonstrating that the colonization of catheters by the fungal pathogen *Candida albicans* can be prevented by coating them with thin polymer multilayers composed of chitosan (CH) and hyaluronic acid (HA) designed to release a β-amino acid-based peptidomimetic of antimicrobial peptides (AMPs). We demonstrate here that this β-peptide is also potent against *S. aureus* (MIC = 4 µg/mL) and characterize its selectivity toward *S. aureus* biofilms. We demonstrate further that β-peptide-containing CH/HA thin-films can be fabricated on the surfaces of rough planar titanium substrates in ways that allow mammalian cell attachment and permit the long-term release of β-peptide. β-Peptide loading on CH/HA thin-films was then adjusted to achieve release of β-peptide quantities that selectively prevent *S. aureus* biofilms on titanium substrates *in vitro* for up to 24 days and remained antimicrobial after being challenged sequentially five times with *S. aureus* inocula, while causing no significant MC3T3-E1 preosteoblast cytotoxicity compared to uncoated and film-coated controls lacking β-peptide. We conclude that these β-peptide-containing films offer a novel and promising localized delivery approach for preventing orthopaedic implant infections. The facile fabrication and loading of β-peptide-containing films reported here provides opportunities for coating other medical devices prone to biofilm-associated infections.

**STATEMENT OF SIGNIFICANCE:** Titanium (Ti) and its alloys are used widely in internal fixation devices due to their mechanical strength and long-term biocompatibility. However, these devices are susceptible to bacterial colonization and the subsequent formation of biofilms. Here we report a chitosan and hyaluronic acid polyelectrolyte multilayer-based approach for the localized delivery of helical, cationic, globally amphiphilic β-peptide mimetics of antimicrobial peptides to inhibit *S. aureus* colonization and biofilm formation. Our results reveal that controlled release of this β-peptide can selectively kill *S. aureus* cells without exhibiting toxicity toward MC3T3-E1 preosteoblast cells. Further development of this polymer-based coating could result in new strategies for preventing orthopaedic implant-related infections, improving outcomes of these titanium implants.

## 1. Introduction

Internal fixation devices (IFDs) are used routinely for the fixation of bone fractures, replacement of arthritic joints, correction and stabilization of the spinal column, and other orthopaedic applications [1]. Titanium and titanium alloys are considered the gold-standard material for IFDs due to their high mechanical stability, low susceptibility to corrosion, inertness, biocompatibility and long-term functionality [1–4]. While the surface topography of titanium is beneficial in the context of promoting osseointegration of the implant, it also promotes microbial colonization [5]. Post-operative microbial infections can occur either within the first two months after implantation and/or many months to years post-surgery and are one of the most common complications following IFD implantation, with infection rates of 1–2.5% for primary knee and hip replacements and up to 20% after revision surgeries have been performed [4,6,7]. Post-operative infections have been linked to aseptic loosening, implant failure, and, in severe cases, morbidity or mortality [3,6,8–15].

*Staphylococci*, including *Staphylococcus aureus* (*S. aureus*), methicillin-resistant *S. aureus* (MRSA), and coagulase-negative *Staphylococci*, are the most common isolated microorganisms from infected IFDs due to their ability to adhere to IFD surfaces and subsequently form biofilms on implants [16,17]. These biofilms can then result in septic arthritis and osteomyelitis [3,7,12,18– 20]. Current treatments for *S. aureus* IFD-associated infections consist of systemic delivery of antibiotics in combination with surgical site radical debridement and/or implant replacements; several revision surgeries are often needed [6,12,14,21,22]. Intravenous antibiotic treatment for the first 2–4 weeks, including rifampicin monotherapy and/or combination therapy with fluoroquinolones, clindamycin and β-lactams followed by an oral antibiotic regimen for an additional 4–6 weeks, remains the standard care for any antibiotic-sensitive *Staphylococcus* infection [20,23]. For the treatment of MRSA infections, combination therapy including vancomycin, dicloxacillin, linezolid, daptomycin and fosfomycin antibiotics is often needed [16,24]. In addition to the severe side effects associated with the systemic delivery of antibiotics and the emergence of resistant *S. aureus* strains, antibiotic treatment for IFD-associated infections faces challenges such as poor antibiotic bioavailability in bone tissue and antibiotic resistance in bacterial biofilms [12,14,21,22]. In view of these challenges, there is a critical need for the development of new strategies preventing microbial infections associated with IFDs.

Antimicrobial peptides (AMPs) and peptidomimetic analogs of AMPs have been studied as potential new classes of antimicrobials. These AMPs are part of the host’s adaptive immune system and often display selective toxicity to microbial cells vs. host cells [25,26]. Structural characteristics such as an overall net positive charge and adopting a global amphiphilic conformation (e.g., α-helix) upon contact with microbial surfaces confer antimicrobial activity [27,28]. The proposed toxicity mechanism of many types of AMPs involves disruption of the microbial cell membrane, leading to membrane permeabilization, cell lysis and subsequent death [25,29]. Given the lack of a single target of AMPs, the development of bacterial resistance to AMPs and their mimetics is thought to be less likely than for traditional antibiotics [26,30].

While AMPs hold promise as antimicrobials, their structural instability under physiologic conditions and susceptibility to protease degradation *in vivo* have limited their development as antimicrobials [31–34]. Recently, β-peptide foldamers have emerged as cationic, globally amphiphilic structurally stable peptidomimetics. β-Peptides have been demonstrated to have strong antibacterial activity against planktonic Gram-positive and Gram-negative bacteria [28,29,35–39]. Motivated by the potential of β-peptides as an alternative antimicrobial treatment, we have previously demonstrated 14-helical β-peptide toxicity toward planktonic *Candida albicans* cells and prevention of *C. albicans* biofilms *in vitro* [28,37,40,41]. Our previous work also identified the (ACHC-β^3^hVal-β^3^hLys)_3_ β-peptide (Scheme 1) as a promising antimicrobial candidate due to its strong activity against planktonic *C. albicans* cells (MIC = 4 µg/mL), ability to prevent *C. albicans* biofilm formation (MIC = 8 µg/ml) and good selectivity to microbial cells (2.3 ± 0.7% hemolysis at planktonic MIC) [37,40]. Here we build upon our previous work to evaluate the ability of (ACHC-β^3^hVal-β^3^hLys)_3_ β-peptide to inhibit the formation of *S. aureus* biofilms and selectivity to *S. aureus* vs. preosteoblast cells

**Scheme 1:**
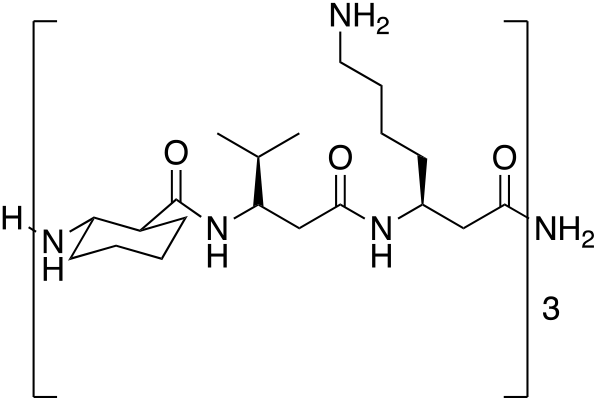
Chemical structure of 14-helical (ACHC-β^3^hVal-β^3^hLys)_3_ β-peptide used in this study.

In an effort to improve the selectivity of AMPs and AMP mimetics, increase activity against microbial cells and reduce the toxicity associated with a systemic delivery, strategies that can localize their delivery to sites prone to infection have recently emerged as an alternative treatment for IFD-related infections. Polyelectrolyte multilayer (PEM) coatings have been developed as a localized platform for surface-mediated release of active biological agents such as growth factors (e.g., BMP-2, bFGF), β-peptides, antibiotics, DNA, among other active agents [22,42–45]. We have also recently reported the prevention of *C. albicans* colonization and biofilm formation *in vitro* and *in vivo* on catheters coated with either polyglutamic acid / poly-L-lysine (PGA/PLL) or chitosan /hyaluronic acid (CH/HA) PEM films loaded with β-peptide [43,46,47]. Motivated by that past work, we focus here on demonstrating the potential use of β-peptide-containing PEM coatings fabricated on the surfaces of rough titanium substrate surfaces for preventing *S. aureus*-related infections. We further evaluated the biocompatibility of these coatings with model osteogenic mammalian cells (MC3T3-E1 preosteoblast). Our results suggest that the controlled release of β-peptide quantities selective only to microbial cells can be achieved using minor modifications, such as chemical crosslinking and by tuning the β-peptide loading. Specifically, our results reveal that β-peptide-containing films deposited on titanium substrates surfaces can release sufficient quantities of β-peptide to prevent *S. aureus* biofilm formation *in vitro* for up to 24 days and five bacterial challenges. Overall, the results reported here indicate β-peptide-containing PEMs coatings to be a useful platform for the design of antibacterial coated IFDs for inhibiting *S. aureus* biofilm formation.

## 2. Materials and Methods

### 2.1. Materials

Branched polyethyleneimine (BPEI, MW=25,000), chitosan (CH, medium molecular weight), phosphate-buffered saline (PBS), paraformaldehyde, glutaraldehyde, menadione, filtered water for cell culture, fluorescein-labeled hyaluronic acid, chloramphenicol, and medical grade titanium disks were purchased from Sigma-Aldrich. Sodium hyaluronate (HA, MW 1,5000,000–2,200,000) was purchased from Acros Organic. 2,3-Bis-(2-methoxy-4-nitro-5-sulfophenyl)-2H-tetrazolium-5-carboxanilide (XTT), RPMI 1640 powder containing L-glutamine and phenol red (without HEPES or Na bicarbonate), penicillin-streptomycin (10,000 U/mL), NaCl, 3-(N-morpholino)propanesulfonic acid (MOPS), 1-ethyl-3-(3-dimethylaminopropyl)carbodiimide hydrochloride (EDC), N-hydroxysulfosuccinimide (Sulfo-NHS), Calcein-AM, ethidium homodimer-1, Hoechst 33342, Nunc™ Lab-Tek™ II Chamber Slide™ System, and Pierce™ Quantitative Fluorometric Peptide Assay were purchased from Thermo Fisher Scientific. α-MEM (1x) minus ascorbic acid was obtained from Gibco. Osmium tetroxide (4%) was obtained from Electron Microscopy Sciences. Accutase was purchased from Innovative Cell Technologies. Cell Titer Glo 2.0 assay kits were obtained from Promega. All materials were used as received.

### 2.2. General Considerations

β-Peptide (ACHC-β^3^hVal-β^3^hLys)_3_ was synthesized using previously reported methods [37]. Titanium substrates were cut to 0.6 cm width x 1.8 cm length dimensions, cleaned with acetone, ethanol, methanol, and deionized water, dried under a stream of filtered and compressed air, and plasma etched for 1800s (Plasma Etch, Carson City, NV) prior to the fabrication of PEM films. Uncoated, film-coated and β-peptide-loaded titanium substrates were UV sterilized for 15 min per side using a biosafety cabinet prior to biological experiments. Fluorescence microscopy images were obtained with an Olympus IX70 epifluorescence microscope using Nikon NIS image acquisition software. Fiji Image J was used to create merged images and quantify fluorescence intensities. Critical point drying, sputtering and scanning electron microscopy (SEM) were performed using a Leica EM CPD 300 critical point dryer, Leica ACE600 Sputter, and a LEO SEM microscope at 5kV. Fluorescence measurements to characterize β-peptide release, absorbance measurements to quantify *S. aureus* cell viability, and luminescence measurements to quantify MC3T3-E1 cell viability were taken with a Tecan M200 multi-well plate reader.

### 2.3. Characterization of β-peptide antimicrobial minimum inhibitory concentration (MIC) and MC3T3-E1 preosteoblast cell cytotoxicity

The antimicrobial activity of (ACHC-β^3^hVal-β^3^hLys)_3_ β-peptide against *S. aureus* biofilms was assayed in 96-well plates according to susceptibility testing guidelines provided by the Clinical and Laboratory Standards Institute [48]. The broth microdilution assay methods were modified to include the quantitative assessment of cell viability using XTT. An aliquot of 100 µL of two-fold serial dilutions of β-peptide in MH medium + 0.5% glucose was mixed with 100 µL of *S. aureus* (grown in TSB medium overnight at 37°C and concentration adjusted to 10^6^ CFU/mL) cell suspension and the plates were incubated for 24 hr at 37°C to allow biofilm formation. Wells lacking β-peptide and wells lacking cells and β-peptide were included as controls. After 24 hr, 100 µL of XTT solution (0.5 g/L in PBS, pH 7.4, containing 3 µM menadione in acetone) was added to all wells, and plates were incubated at 37°C in the dark for 1 hr. The supernatants (75 µL) were transferred to a new 96-well plate and absorbance at 490 nm was recorded. The cell viability was normalized to the untreated control and plotted as a function of β-peptide concentration. The lowest assayed concentration of β-peptide that resulted in a decrease in absorbance of at least 90% of the mean was determined to be the minimum inhibitory concentration (MIC) of the peptide.

MC3T3-E1 cell toxicity was assessed using the Cell Titer Glo protocol to quantify the amount of ATP present in metabolically active cells. MC3T3-E1 cells were cultured in MEMα medium supplemented with 10% FBS, 1% of penicillin-streptomycin, in a humidified incubator at 37°C and 5% CO_2_ in air. Upon reaching 100% confluency, cells were detached using Accutase, centrifuged and resuspended in MEMα medium to a final concentration of 5×10^4^ cells/cm^2^. An aliquot of 100 µL of two-fold serial dilutions of β-peptide in MEMα medium was mixed with 100 µL of the MC3T3-E1 cell suspension and the plates were incubated for 24 hr at 37°C and 5% CO_2_ in air. Wells lacking β-peptide and wells lacking cells and β-peptide were included as controls. Afterwards, Cell Titer Glo reagent (100 µL) was added into each well, incubated for 5 min, and the luminescence signal was recorded. The percent of cell death in each well was calculated as:

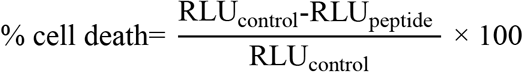

where RLU_control_ represents the luminescence signal of untreated control (well lacking β-peptide) and RLU_peptide_ represents the luminescence signal of β-peptide-containing samples. The percent of cell death was plotted as a function of β-peptide concentration to generate the dose-response curve for MC3T3-E1 toxicity. The IC_20_ value was determined as the β-peptide concentration that resulted in 20% death of MC3T3-E1 cells.

### 2.4. Fabrication of polyelectrolyte multilayers films on the surfaces of titanium substrates

Solutions of HA and BPEI (1 mg/mL) were prepared in deionized water containing 0.15 M NaCl. CH solution (1 mg/mL) was prepared in 0.1 M acetic acid and deionized water containing 0.15 M NaCl. PEMs were fabricated on the surfaces of cut, cleaned, and plasma etched titanium substrates using the following general protocol: (1) substrates were submerged in the 1 mg/mL BPEI solution for 30 min, (2) substrates were removed and immersed in a rinse bath of deionized water containing 0.15 M NaCl for 1 min, followed by a second rinse bath for 1 min, (3) substrates were immersed in the 1 mg/mL CH solution for 5 min, (4) substrates were removed and rinsed as described in step 2, (5) substrates were immersed in the 1 mg/mL HA solution for 5 min, (6) substrates were removed and rinsed as described in step 2 and steps 3–6 were repeated until a total of 19.5 bilayers were deposited. For experiments designed to characterize film growth profiles, PEMs were fabricated as described above but using fluorescein-labeled hyaluronic acid. Fluorescence images from 3 different regions of the PEM-coated titanium substrate were taken after 4.5, 9.5, 14.5 and 19.5 bilayers were deposited. The fluorescence intensities of these images were quantified using Fiji Image J software.

### 2.5. Chemical crosslinking of PEM films deposited on titanium substrates

CH/HA PEMs films were chemically crosslinked by immersing PEM-coated titanium substrates in a 400 mM EDC/100 mM Sulfo-NHS solution in deionized water containing 0.15 M NaCl for 16 hr at room temperature. Next, substrates were rinsed 3 times for 30 min each in fresh deionized water containing 0.15 M NaCl, followed by a drying using filtered and compressed air. An uncoated control titanium substrate was immersed in deionized water containing 0.15 M NaCl without crosslinking agents for 16 hrs as control. For epifluorescence microcopy to characterize the films before and after crosslinking, CH/HA films were fabricated using fluorescein-labeled HA and fluorescence images were taken before and after crosslinking.

### 2.6. Characterization of crosslinked films using PM-IRRAS

To characterize CH/HA PEM film crosslinking, CH/HA PEMs films were deposited on gold-coated silicon substrates and crosslinked as described above. Crosslinking was characterized by polarization-modulation infrared reflectance-absorbance spectroscopy (PM-IRRAS) conducted in a similar fashion to previously reported methods [49]. Briefly, gold-coated silicon substrates coated with CH/HA PEM films before and after crosslinking were placed in a Nicolet Magna-IR 860 Fourier transform infrared spectrophotometer equipped with a photoelastic modulator (PEM-90, Hinds Instruments, Hillsboro, OR), a synchronous sampling demodulator (SSD-100, GWC Technologies, Madison WI), and a liquid nitrogen cooled mercury-cadmium-telluride detector. The modulation was set at 5865.0 nm, 0.5 retardation and 500 scans with a resolution of 2 cm^-1^ were obtained for each sample. The differential reflectance infrared spectra were normalized and converted to absorbance spectra using a previously reported procedure [49].

### 2.7. PEM loading with β peptide

Titanium substrates coated with crosslinked CH/HA PEMs were immersed in a 0.44 mg/mL solution (or an otherwise desired concentration) of β-peptide (ACHC-β^3^hVal-β^3^hLys)_3_ in deionized water containing 0.15 M NaCl for a period of 24 hr at room temperature. β-Peptide loaded substrates were removed from solution and dried under a stream of filtered and compressed air. An uncoated control titanium substrate was immersed in deionized water containing 0.15 M NaCl without β-peptide for 24 hrs. Similarly, a PEM film-coated control was immersed in in deionized water containing 0.15 M NaCl without β-peptide for 24 hr.

### 2.8. Estimation of film thickness

The thicknesses of CH/HA, crosslinked CH/HA, and β-peptide-loaded CH/HA films on titanium substrates were estimated using focused ion beam scanning electron microscopy (FIB-SEM). β-Peptide loaded CH/HA films were prepared as described before. Uncrosslinked CH/HA films were incubated for a total of 30 hrs in deionized water containing 0.15 M NaCl as a control lacking crosslinking solution and β-peptide. Similarly, control crosslinked CH/HA films were incubated in deionized water containing 0.15 M NaCl and lacking β-peptide for 24hrs. Samples were platinum-palladium coated and 1 rectangular section 10 µm wide was milled using a 0.10 nA electrical current to create a film cross-section. Five different regions within the milled section were selected to estimate the film thickness using SEM at 2kV.

### 2.9. Characterization β-peptide release from PEM films

Characterization of β-peptide release from PEMs on titanium substrates was performed by following the manufacturer’s specifications for the Pierce™ Quantitative Fluorometric Peptide Assay kit. Briefly, uncoated, film-coated, and β-peptide loaded titanium substrates were immersed in 750 µL of filtered water and incubated at 37°C. At predetermined intervals, titanium substrates were removed from the incubator and β-peptide concentration in the release solution was quantified. 10 µL of the release solution was mixed with 70 µL of peptide assay buffer and 20 µL of peptide assay reagent and incubated for 5 min. Fluorescence intensity was recorded at 390 nm excitation and 475 nm emission. Fluorescence measurements were converted to β-peptide concentrations using a calibration curve constructed with known β-peptide concentrations. After each measurement, titanium substrates were immersed in 750 µL of fresh filtered water and returned to the incubator. The plot shown in Figure 4 was constructed by cumulatively adding the concentrations of β-peptide released at each timepoint and is normalized to the titanium substrate surface area.

### 2.10. Characterization of the antibacterial activity of β-peptide-loaded PEM films

*S. aureus* ATCC 3359 and AH17456 cells were grown overnight at 37°C in liquid TSB, subcultured the following day, and grown to an optical density at 600 nm (OD^600^) of 0.4. TSB growth medium for the GFP-expressing AH1756 strain was supplemented with 10 µg/mL chloramphenicol for plasmid maintenance purposes. Cells were washed with PBS and resuspended in MH medium supplemented with 0.5% glucose to a cell density of 10^6^ CFU/mL to stimulate biofilm growth. Uncoated, film-coated, and β-peptide loaded substrates were placed inside a four-well Lab Tek chamber containing 750 µL of *S. aureus* cell suspension in supplemented MH medium and incubated for 24 hr at 37°C to allow biofilm growth. Growth of biofilms was characterized using (i) an XTT metabolic activity assay and (ii) by imaging the biofilms using fluorescence microscopy and SEM.

For the XTT metabolic activity assay, each titanium substrate was removed from the four-well Lab Tek chamber, gently washed with PBS and transferred into a new and unused Lab Tek chamber. XTT solution (750 µL; 0.5 g/L in PBS, supplemented with 3 µM menadione in acetone) was added to each well of the Lab Tek Chamber containing the uncoated, film-coated and β-peptide loaded titanium substrates. After incubating the XTT solution at 37°C for 1.5 hr in the dark, 75 µL of the supernatant was transferred into a 96-well plate and the absorbance of the solution at 490 nm was measured to determine the relative metabolic activity of the biofilms. Data were plotted relative to the absorbance value from the well containing the uncoated titanium substrate control.

A biological and XTT metabolic activity assay configuration similar to that described above was used to evaluate biofilm formation after multiple *S. aureus* challenge experiments and after incubation of substrates in PBS prior to biofilm formation. For the multiple challenge experiments, uncoated, film-coated, and β-peptide loaded titanium substrates were initially incubated with an *S. aureus* inoculum and biofilms were allowed to grow for 24 hr (challenge 1). Substrates were then incubated in PBS for an additional 2 days and subsequently challenged with an additional *S. aureus* inoculum for 24 hr (challenge 2). This multiple-challenge process was repeated until 5 different *S. aureus* challenges were achieved, for a total of 18 days. Different sets of titanium substrates were sacrificed after 1, 2, 3, 4, and 5 challenges and XTT metabolic assays were performed to quantify biofilm formation. For the PBS pre-incubation experiments, uncoated, film-coated, and β-peptide loaded titanium substrates were incubated in PBS at 37°C for the specified period of time (e.g., 1, 2, 4, 6, 12, 24, 36, 48, and 60 days) and then challenged with an *S. aureus* inoculum. Extents of biofilm formation were quantified via XTT assay and data were normalized to the uncoated control.

To analyze biofilm formation using fluorescence microscopy, a GFP-expressing *S. aureus* strain AH1756 was used. Following 24 hr biofilm formation at 37°C, uncoated, film-coated, and β-peptide loaded titanium substrates were washed with PBS and biofilm growth was inspected under an epifluorescence microscope. We also evaluated *S. aureus* biofilm morphology using SEM. For this analysis, uncoated, film-coated, and β-peptide loaded titanium substrates were prepared using a previously published protocol [43,46]. Briefly, titanium substrates were placed in a fixative solution (1% (v/v) glutaraldehyde and 4% (v/v) paraformaldehyde) overnight at 4 °C. Next, titanium substrates were rinsed with PBS for 10 min and placed in osmium tetroxide (1%) for 30 min, followed by another wash in PBS for 10 min. Titanium substrates were then dehydrated using a series of ethanol washes (30%, 50%, 70%, 85%, 95% and 100%, 10 min each) and final desiccation was performed using critical point drying. Finally, specimens were mounted in aluminum stubs and sputter-coated with a 12 µm thick layer of platinum-palladium. Samples were then imaged by SEM in high-vacuum mode at 5kV.

### 2.11. Biocompatibility of β-peptide-loaded PEM films

MC3T3-E1 cells were grown in MEMα medium, supplemented with 10% FBS and 1% penicillin-streptomycin, in a humidified incubator at 37°C and 5% CO_2_ in air. Culture medium was replenished every 2–3 days and cells were sub-cultured using Accutase when near-100% confluency was observed. All cells used in these studies were less than passage number 25. Uncoated, film-coated, and β-peptide loaded titanium substrates were placed inside four-well Lab Tek chambers containing 750 µL of an MC3T3-E1 cell suspension adjusted to a cell density of 5×10^4^ cells/cm^2^ in α-MEM and incubated for 24 hr at 37°C and 5% CO_2_, unless otherwise specified. MC3T3-E1 cell viability was characterized using (i) a Cell Titer Glo metabolic activity assay and (ii) by visualizing MC3T3-E1 cell attachment with fluorescence microscopy.

Cell Titer Glo assessment of MC3T3-E1 cell metabolic activity was performed according to the manufacturer’s recommendations. Briefly, uncoated, film-coated, and β-peptide loaded titanium substrates were removed from the four-well Lab Tek chambers, gently washed with PBS and transferred into a new and unused Lab Tek chamber. MEMα medium and Cell Titer Glo reagent (750 µL each) were added into the wells and incubated for 5 min. Next, 180 µL of supernatant was transferred into a 96-well plate and luminescence signal was quantified. Background luminescence from wells containing medium and Cell Titer Glo was subtracted from all readings and data were normalized relative to the uncoated titanium control. For the PBS pre-incubation experiments, uncoated, film-coated, and β-peptide-loaded titanium substrates were allowed to elute β-peptide in PBS at 37°C for the specified period of time (e.g. 1, 2, 4, 6, 12, 24, 36, 48 and 60 days). MC3T3-E1 cells were then seeded on the films and allowed to attach and grow for 24 hr. Viability of the MC3T3-E1 cells was then quantified using the Cell Titer Glo assay.

Fluorescence microscopy images of MC3T3-E1 cells on the surfaces of uncoated, film-coated, and β-peptide-loaded titanium substrates were acquired as previously described [22]. Briefly, each titanium substrate was placed inside a four-well Lab Tek chamber containing 750 µL of a MC3T3-E1 cell suspension adjusted to a cell density of 5×10^4^ cells/cm^2^ in MEMα medium and incubated for the specified period of time (e.g. 2, 4, and 6 days) at 37°C and 5% CO_2_. Afterwards, a working solution of 2 µM Calcein-AM, 4 µM ethidium homodimer-1 and 0.2 mg/mL Hoechst 33342 was prepared in PBS and then incubated with the cells on Ti substrates for 30 min at 37°C. The dye solution was gently aspirated from the wells, and then substrates were rinsed with PBS and imaged using an epifluorescence microscope.

### 2.12. Statistical analysis

GraphPad Prism 7.0 (GraphPad Software, Inc) was used for all statistical analysis. For pairwise comparisons a Student’s T-test was performed. Statistical comparisons were performed using two-way analysis of variance (ANOVA) or one-way ANOVA as appropriate, with Tukey’s Honest Significant Difference post-hoc analysis for multiple testing over all comparisons. Figure legends describe the statistical tests used for each particular data set. Statistical significance was accepted at a p value of less than 0.05. Data are represented as mean values ± standard deviations (SD) for three separate biological replicates, with three technical replicates in each biological replicate.

## 3. Results and Discussion

### 3.1. β-peptide inhibits *S. aureus* biofilm formation

The ability of *S. aureus* cells to attach and form drug-resistant biofilms on the surfaces of IFDs poses a challenge for the treatment of implant-related bacterial infections. Motivated by previous studies demonstrating the antibacterial activity of cationic 14-helical β-peptides, with MICs against planktonic *S. aureus* cells ranging from 3.1 to 200 µg/mL [29,35,50], and previous work demonstrating (ACHC-β^3^hVal-β^3^hLys)_3_ β-peptide activity and selectivity against *C. albicans* cells vs. human red blood cells [37,40], we tested the potential of the (ACHC-β^3^hVal-β^3^hLys)_3_ β-peptide to prevent the formation of *S. aureus* biofilms. We quantified the MIC for preventing *S. aureus* biofilms in 96-well polystyrene plates, following the CLSI antimicrobial susceptibility standards [48], with modifications to include biofilm growth (e.g. 37°C, MH medium supplemented with 0.5% glucose and 24 hr incubation time). Results shown in Figure 1 (orange squares) show that (ACHC-β^3^hVal-β^3^hLys)_3_ β-peptide has a biofilm inhibition MIC of 4 μg/mL. Although strong antimicrobial activity is highly desired, evaluating selectivity to *S. aureus* cells exclusively is also crucial for potential use to prevent IFD-related infections *in vivo*. We therefore, investigated (ACHC-β^3^hVal-β^3^hLys)_3_ β-peptide biocompatibility with MC3T3-E1 preosteoblast subclone 4 cells, a model mammalian cell line with osteoblast differentiation capacity and mineralization activity. Our results showed a concentration-dependent β-peptide toxicity toward MC3T3-E1 preosteoblast cells with an inhibitory concentration resulting in 20% cell death (IC_20_) of 22.6 ± 7.4 μg/mL (Figure 1, blue circles). We defined an *in vitro* selectivity index (SI) as the ratio of β-peptide cytotoxicity (IC_20_) against MC3T3-E1 cells to MIC against inhibiting *S. aureus* biofilm formation (SI= IC_20_/MIC). Using this approach, (ACHC-β^3^hVal-β^3^hLys)_3_ β-peptide was demonstrated to have a SI value of 5.7, suggesting good selectivity for *S. aureus* vs. MC3T3-E1 cells. This result motivated the development of delivery strategies for its localized release for preventing *S. aureus* biofilm formation.

**Figure 1:**
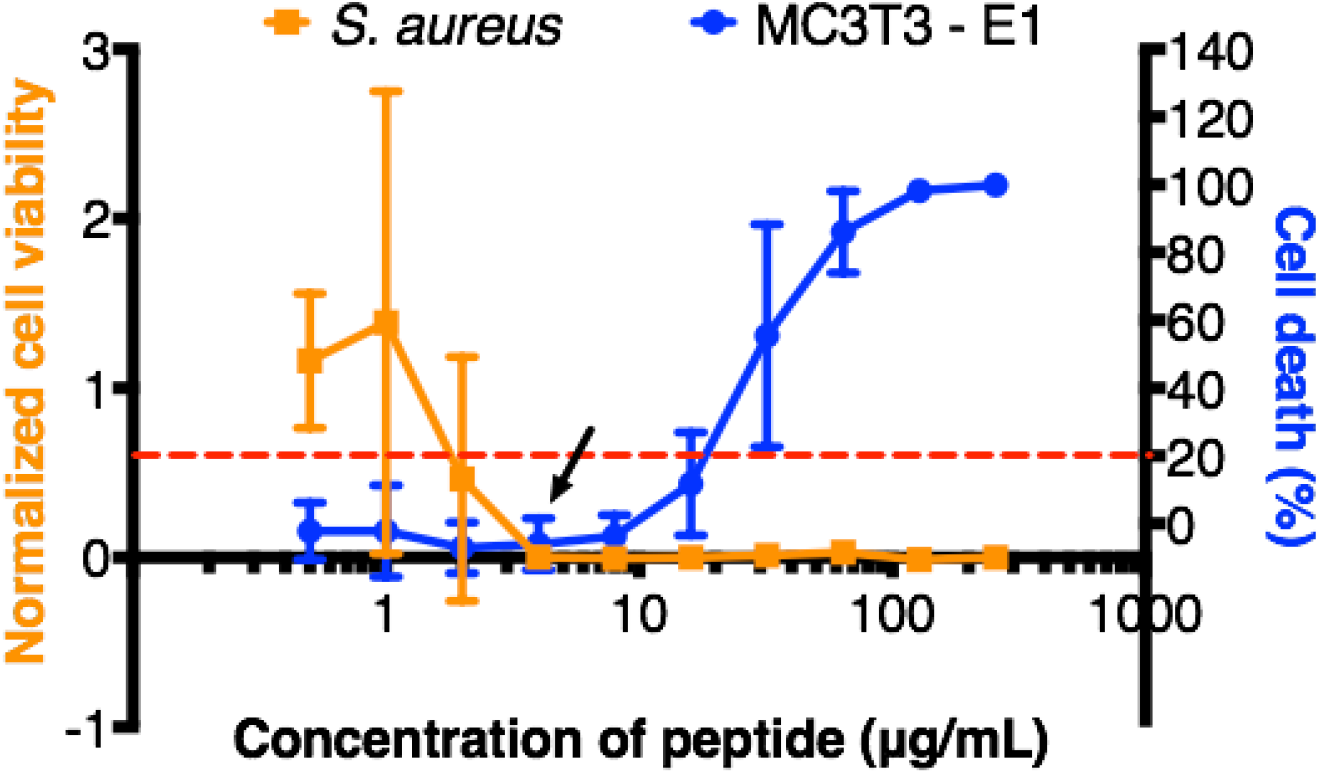
Effect of β-peptide concentration on inhibition of *S. aureus* biofilms and toxicity toward MC3T3-E1 cells. *S. aureus* cells (10^6^ cells/mL) were incubated with the indicated β-peptide concentrations in MH medium + 0.5% glucose in 96-well plates for 24 hr at 37°C, and then biofilm was quantified using an XTT assay. Biofilm viability was normalized to a control lacking β-peptide. The arrow indicates the MIC for inhibiting *S. aureus* biofilm formation. To evaluate β-peptide cytotoxicity, we incubated MC3T3-E1cells (5×10^4^ cells/cm^2^) with the indicated β-peptide concentrations in MEM α medium in 96-well plates for 24 hr at 37°C and 5% CO_2_. MC3T3-E1 cell viability was quantified using a Cell Titer Glo assay. Cell death was calculated based on the percent change with respect to the cells grown in the absence of peptide. The dashed red line indicates the IC_20_ β-peptide concentration. Data points represent the mean values and error bars the standard deviation of three independent experiments.

### 3.2. Fabrication and characterization of β-peptide-containing PEM films

We selected the polysaccharide-based CH/HA PEM film system for use in this study because these coatings have been well-studied as a platform for the localized release of active agents from antimicrobial coatings and tissue-integrating scaffolds [43,51–55]. Additionally we have recently demonstrated antifungal activity of CH/HA PEMs containing β-peptide [43,47]. That study showed that CH/HA PEM films containing (ACHC-β^3^hVal-β^3^hLys)_3_ fabricated in the lumens of catheter segments using an iterative flow-based approach prevented *C. albicans* biofilms *in vitro* and *in vivo*. This current study sought to extend upon that prior work to (i) determine whether CH/HA PEMs fabricated on rough and planar titanium substrates could be used to promote the long-term release of β-peptide and prevent formation of *S. aureus*-related biofilms, (ii) evaluate the biocompatibility of β-peptide-loaded PEM films with the MC3T3-E1 preosteoblast cell line, and (iii) assess the selectivity of these β-peptide-containing coatings against *S. aureus* vs. MC3T3-E1 cells.

Using an iterative immersion-based layer-by-layer assembly approach, we deposited CH/HA multilayers on the surfaces of titanium substrates (Figure 2) and characterized their growth by monitoring the fluorescence intensity of FITC-labeled HA incorporated within the films, using a previously published approach consisting of imaging and analyzing the fluorescence intensity after the deposition of every five CH/HA bilayers [46]. As shown in Figure 2, the average fluorescence intensity increased with the number of CH/HA bilayers deposited, consistent with layer-by-layer film growth [56].

**Figure 2:**
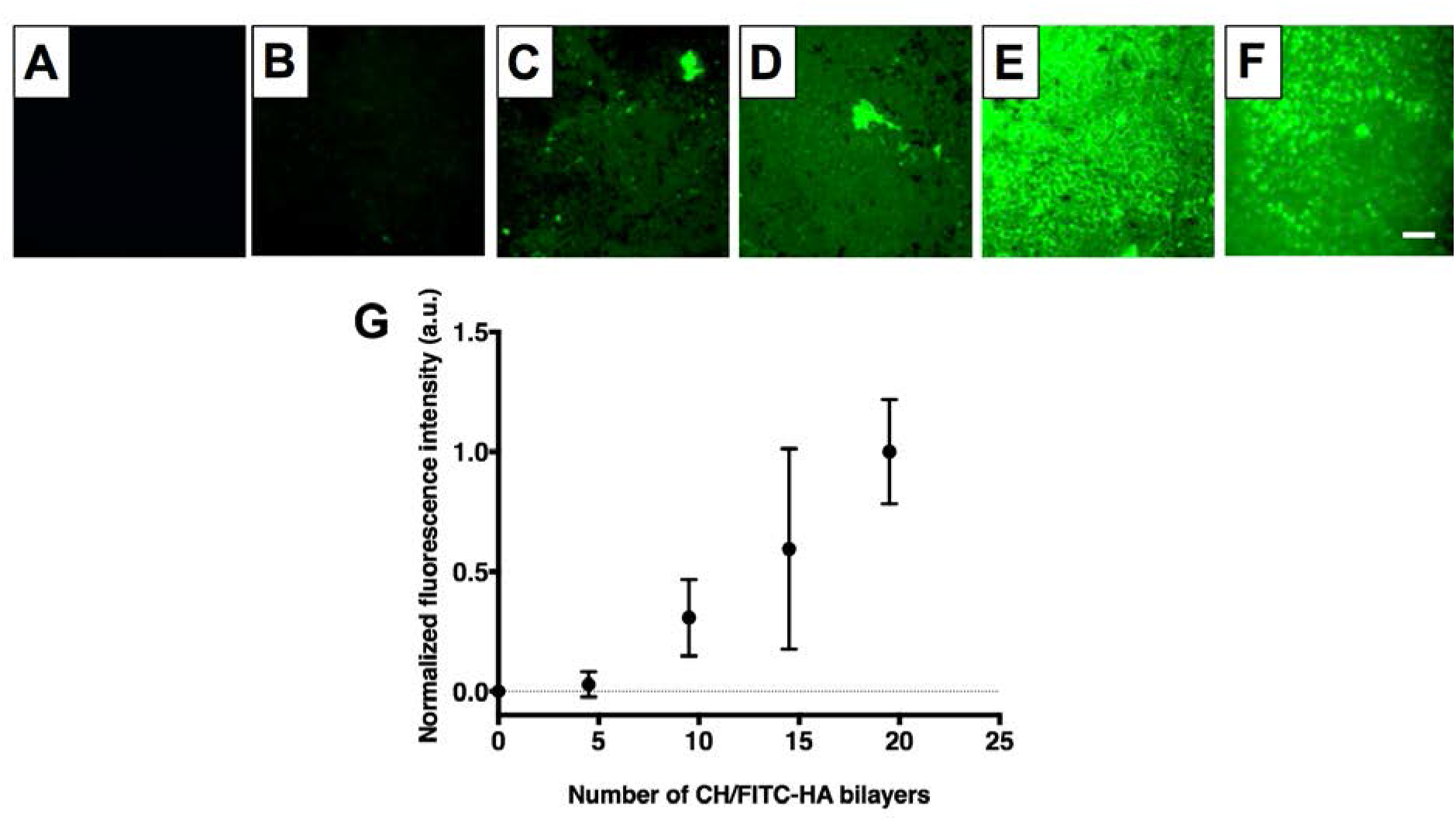
CH/HA film deposition on titanium substrates. A-E) Representative epifluorescence images of (CH/FITC-HA)x coated titanium substrates; where x= 0 (A), 4.5 (B), 9.5 (C), 14.5 (D), 19.5 (E). (F) Representative epifluorescence image of (CH/FITC-HA)_19.5_ film deposited on titanium after EDC/NHS cross-linking. G) Growth profile of crosslinked CH/HA films. Fluorescence intensity was quantified using Fiji Image J. Data points are the average mean values and error bars are the standard deviation of at least three regions of each titanium substrate corresponding to three independent experiments, normalized to the fluorescence intensity of 19.5 CH/FITC-HA bilayers. Scale bar: 260 µm.

We then characterized the ability of MC3T3-E1 preosteoblast cells to attach and proliferate on CH/HA film-coated titanium substrates over a period of 6 days. Visual inspection of cell attachment and quantification of proliferation with a Cell Titer Glo assay revealed the extent of MC3T3-E1 cells attachment on CH/HA film-coated titanium substrates (Figure 3 D-F, S4) was no different than on uncoated titanium (Figure 3 A-C, S4). However, proliferation over a period of 6 days on CH/HA film-coated titanium substrates (Figure 3 D-F, I, orange bars) was lower than on uncoated titanium (Figure 3 A-C, I, blue bars), suggesting that the surfaces of the film-coated substrates were less favorable for supporting MC3T3-E1 cell growth than bare titanium. Cellular adhesion and proliferation on PEM-coated surfaces is known to depend upon film physical and mechanical properties (e.g. Young’s modulus, roughness, and degree of hydration) [55–57]. For example, previous studies have demonstrated that depositing poly(allylamine hydrochloride – poly(acrylic acid) films at a higher pH resulted in more rigid films that increased the adhesion of NR6WT fibroblasts [58]. In addition, Schneider *et. al*. demonstrated that chemical crosslinking of CH/HA films increased film roughness and rigidity and enhanced the viability of attached HT29 cells [59].

As part of a strategy to improve MC3T3-E1 cell proliferation on our CH/HA films we chemically crosslinked the films using an EDC/NHS (400 mM EDC/100 mM NHS in 0.15 M NaCl) carbodiimide treatment in a post-fabrication step [60–62]. This carbodiimide-based crosslinking catalyzes the formation of amide bonds between the carboxylic groups of HA and the amine groups of CH, and was selected because both crosslinking agents are water soluble and carbodiimides do not remain as part of the amide linkage, but instead are converted to nontoxic, water soluble urea derivatives that can be easily removed [63,64]. Following the crosslinking of CH/HA films containing FITC-labeled HA, we characterized the films by fluorescence microscopy (Figure 2 E-F). We also included CH/HA films incubated in deionized water containing 0.15 M NaCl during the crosslinking step as an uncrosslinked control (Figure S2). Our results demonstrate that, after crosslinking, the surface remained covered by the films (Figure 2F, S2B), similar to control CH/FITC-HA films that were incubated in deionized water containing 0.15 M NaCl (Figure S2C). We confirmed CH/HA film crosslinking using polarization modulation infrared reflection-absorption spectroscopy (PM-IRRAS). Because PM-IRRAS requires a reflective surface, we deposited the CH/HA PEMs on gold-coated silicon wafers and obtained the infrared spectra of the films before and after EDC/NHS crosslinking. Inspection of the spectra revealed disappearance of the HA carbonyl group at 1412 cm^-1^ after film crosslinking (Figure S3), consistent with the formation of an amide bond between CH and HA. Additionally, we observed that the crosslinked films had a strong absorbance in the amide I (1660 cm^-1^) and amide II bands (1570 cm^-1^), further suggesting the successful crosslinking of the CH/HA films (Figure S3). We then compared MC3T3-E1 cell adhesion and proliferation on crosslinked and uncrosslinked films. We did not observe a significant change in MC3T3-E1 cell adhesion on the surfaces of the uncoated substrates, uncrosslinked films, and crosslinked films (Figure 3 and S4). However, CH/HA film crosslinking enhanced cell MC3T3-E1 cell proliferation over 6 days compared to uncrosslinked films, leading to a similar number of cells as the bare titanium surface (Figure 3). These results demonstrate that film crosslinking leads to coatings that can support MC3T3-E1 cell viability and proliferation, likely by modulating film rigidity, roughness, and/or degree of hydration as demonstrated in past studies [55–57,59].

**Figure 3:**
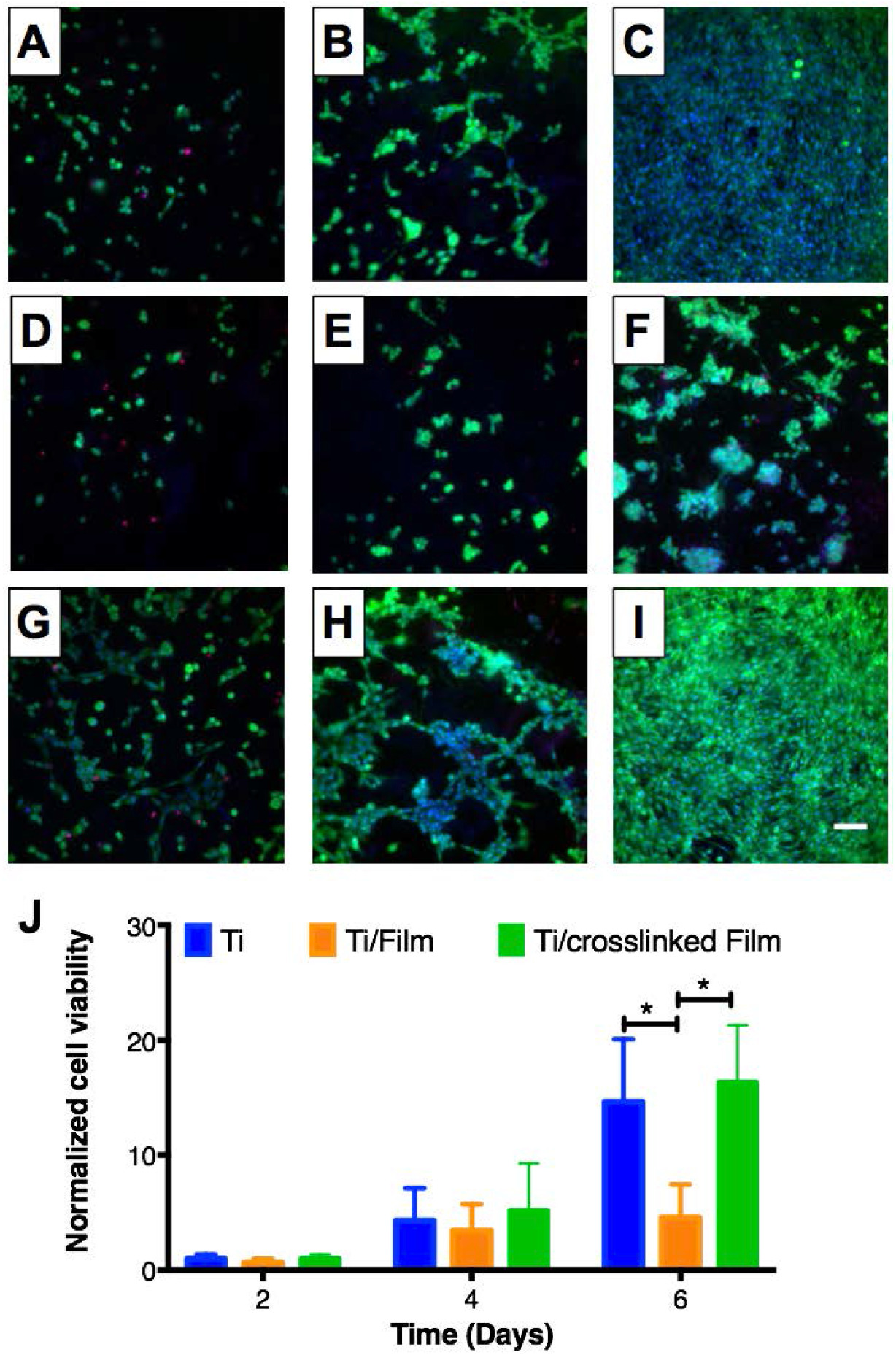
MC3T3-E1 cell attachment and proliferation on titanium substrates coated with 19.5 bilayer thick CH/HA films. A-I) Fluorescent micrographs of live cells (Calcein AM, green), dead cells (propidium iodide, red) and nuclei (Hoechst, blue) of representative fields of (A-C) uncoated, (D-F) CH/HA film-coated and (G-I) crosslinked CH/HA film-coated titanium substrates at day 2 (A, D, and G), day 4 (B, E, and H) and day 6 (C, F, and I). Scale bar: 175 µm. J) Plot showing quantification of MC3T3-E1 cell viability using the Cell Titer Glo assay as a function of time after seeding on uncoated, CH/HA film-coated, and crosslinked CH/HA film-coated titanium substrates. MC3T3-E1 cell viability is normalized to results obtained using an uncoated control. Data points represent the mean values and error bars are the standard deviation of three independent experiments. Asterisks (*) indicate p < 0.01 by two-Way ANOVA using Tukey’s multiple comparison test.

Following the fabrication of crosslinked CH/HA films on titanium substrates, β-peptide (ACHC-β^3^hVal-β^3^hLys)_3_ was loaded into the films by incubating the films in a 0.15 M NaCl solution containing the β-peptide [43,46,47]. For selective bacterial biofilm prevention without toxicity to mammalian cells, β-peptide must elute to achieve a local concentration toxic to *S. aureus* but nontoxic to MC3T3-E1 preosteoblast cells. Therefore, we investigated the effects of varying the concentration of β-peptide in the loading solution on *S. aureus* and MC3T3-E1 cell viability after being cultured for 24 hr on uncoated titanium, on film-coated substrates lacking β-peptide, and on film-coated substrates loaded with β-peptide (Figure 4A). This approach of reporting β-peptide loading concentration was used due to difficulties associated with accurately quantifying the amount of β-peptide loaded into the films (e.g. by post-loading extraction). Past studies from our group have shown that varying β-peptide concentration in the loading solution leads to consistent and measurable differences in both the amount of β-peptide loaded and the release profiles of β-peptide from the films [43,46,65]. Our results demonstrate that the extents of *S. aureus* biofilm inhibition and toxicity toward MC3T3-E1 cells were dependent upon the concentration of β-peptide in the loading solution (Figure 4A). A β-peptide loading solution concentration of 0.44 mg/mL lead to coatings that completely inhibited *S. aureus* biofilm formation and maintained at least 50% survival of MC3T3-E1 cells at 24 hr (Figure 4A). Thus, this loading concentration was selected for further characterization of antimicrobial activity and biocompatibility of β-peptide-containing films.

We next investigated the kinetics of β-peptide release from crosslinked CH/HA films on titanium substrates by incubating uncoated titanium, films lacking β-peptide, and films loaded with β-peptide in deionized water for predetermined amounts of time. Figure 4B shows the cumulative release profile of β-peptide (ACHC-β^3^hVal-β^3^hLys)_3_ from crosslinked films over a period of 54 days. Our results reveal that β-peptide is released gradually at a constant rate of 4.6 ± 2.2 µg/cm^2^/day over a period of 28 days. Over this period, the crosslinked films released 139.4 ± 20.9 µg/cm^2^ of β-peptide. The release profile reported in this study is different from previously published profiles for the release of β-peptide from CH/HA films fabricated on the inner lumens of catheter tube segments, which eluted approximately 350 µg/mL of β-peptide over a period of 100-150 days [43,47]. We note that, for the purposes of this study achieving β-peptide release quantities that were selective to *S. aureus* vs MC3T3-E1 cells was a primary focus; the β-peptide concentration selected for loading these films was thus significantly lower than that used in our previous studies. In addition, chemical crosslinking of CH/HA films, differences in β-peptide sequence, the changes in underlying substrate properties and film-fabrication protocols (e.g., immersion versus flow-based methods) could also contribute to these differences in loading and release. We note further that the release profile reported here is appropriate in the context of IFDs, because the constant rate of β-peptide release was tuned to achieve β-peptide concentrations near the *S. aureus* biofilm MIC (4 µg/mL, Figure 1) over extended periods of time. Also, the localized release of β-peptide reported here is promising in the context of achieving sub-MIC drug concentrations at specific high risk sites, such as IFDs surfaces, which not only results in enhanced effectiveness against preventing *S. aureus* biofilms but also reduces the possibility of microbial cells developing β-peptide resistance [66]. Finally, by controlling the β-peptide release concentration we also reduce potential β-peptide toxicity against MC3T3-E1 preosteoblast cells, thereby mitigating adverse effects on osseointegration.

Finally, we also evaluated how the film thickness changed upon the chemical crosslinking of these films and upon β-peptide incorporation [43,46]. We used a focused ion beam-scanning electron microscope (FIB-SEM) to generate vertical cross-section images of noncrosslinked films, crosslinked films lacking β-peptide, and crosslinked films loaded with β-peptide. Crosslinked films incubated in β-peptide solution had a thickness of 705 ± 146 nm (Figure 4D, S1), significantly greater than crosslinked films lacking β-peptide (148 ± 90 nm; p < 0.001; Figure 4C, S1). The thickness of noncrosslinked films (224 ± 73 nm) was not significantly different from the thickness of crosslinked films (Figure S1). The increase in film thickness after loading with β-peptide is in accordance with those of previous studies on uncrosslinked CH/HA films fabricated using other methods [43,46].

**Figure 4:**
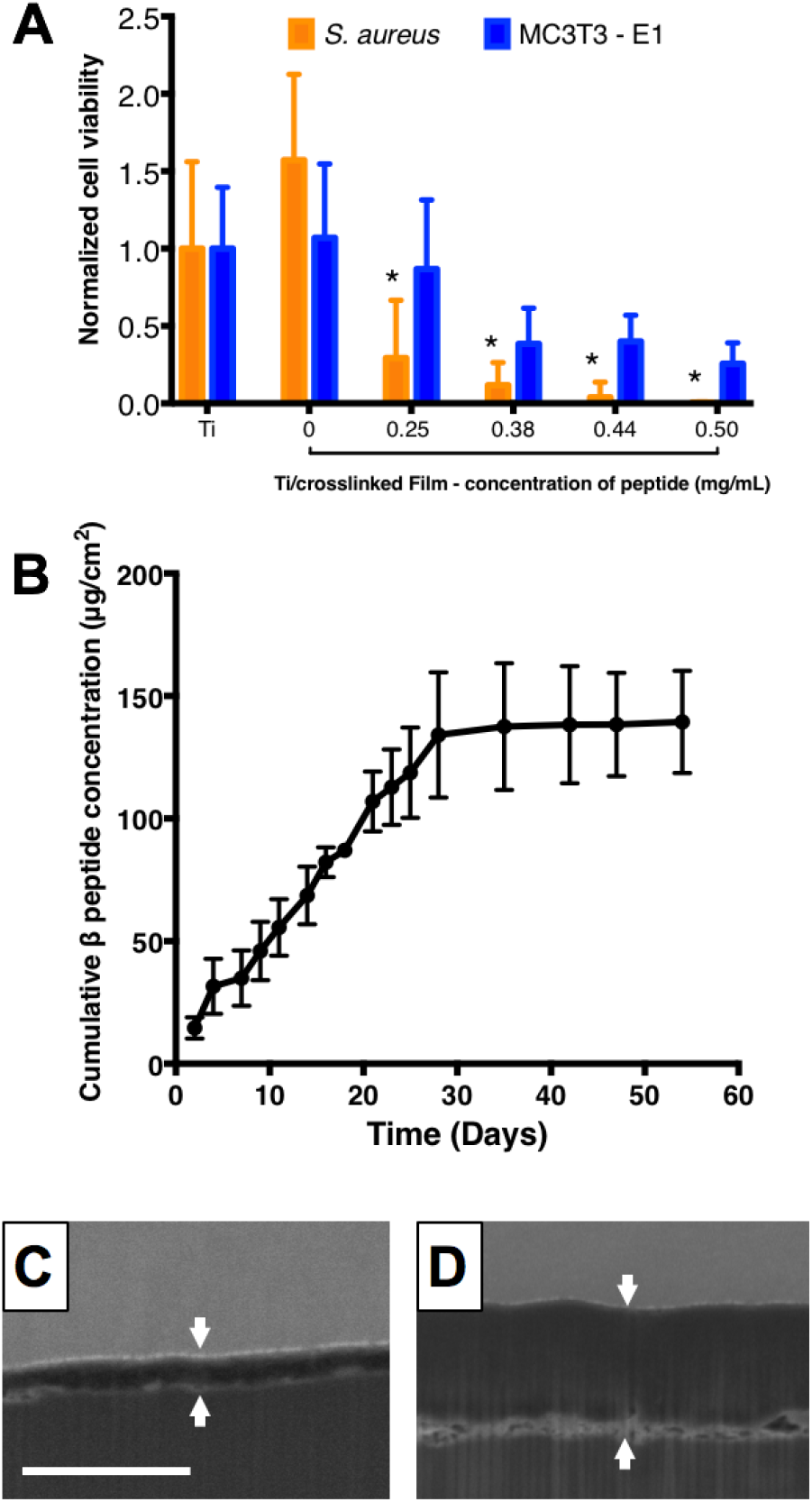
β-peptide loading and release from titanium substrates coated with crosslinked CH/HA films. 19.5 bilayer thick CH/HA films were deposited and crosslinked for 16 hr using an EDC/NHS solution in 0.15 M NaCl. Incorporation of β-peptide was performed by incubating the films for 24 hr in a β-peptide solution in deionized water containing 0.15M NaCl. A) Quantification of *S. aureus* and MC3T3-E1 cell viability on β-peptide loaded CH/HA films as a function of β-peptide loading concentration. After CH/HA film fabrication and β-peptide loading, *S. aureus* or MC3T3-E1 cells were inoculated on the films and allowed to grow for 24 hr. Viabilities were quantified using XTT and Cell Titer Glo assays, respectively. B) Plot showing the cumulative release of β-peptide into PBS (750 µL) at 37°C as a function of time. β-Peptide release concentrations were quantified by using the Pierce quantitative fluorometric assay, calibrated with a standard curve generated with known β-peptide concentrations. C-D) Representative FIB-SEM images of PEM film cross-sections c) before and D) after β-peptide loading. White arrows denote the film edges. Scale bar: 3 µm. Data points represent the mean values and error bars are the standard deviation of three independent experiments.

### 3.3. β peptide-containing coatings inhibit *S. aureus* biofilms

The antimicrobial activity of titanium substrates coated with β-peptide-containing films was characterized by incubating uncoated titanium, coatings without β-peptide, and coatings loaded with β-peptide with an inoculum of 10^6^ *S. aureus* CFU/mL in MH medium supplemented with 0.5% glucose at 37°C for 24 hr. The extent of biofilm formation was then evaluated i) qualitatively by inspecting biofilm density using the GFP-expressing *S. aureus* strain (AH1756) and SEM imaging for characterization of biofilm morphology, and ii) quantitatively by measuring biofilm metabolic activity via an XTT assay. The fluorescence micrographs in Figure 5A-C reveal that robust biofilms formed on the surfaces of uncoated titanium and film-coated titanium substrates without β-peptide, but that β-peptide-loaded films completely inhibited biofilm formation. SEM images reveal biofilms on uncoated substrates and substrates coated with control films without β-peptide to be composed of spherical bacterial 3D cell clusters encased by matrix (Figure 5D-E and Figure S5). In contrast, biofilm was not observed on β-peptide-loaded films by SEM imaging (Figure 5F and Figure S5), in accordance with the fluorescence micrographs. Finally, quantification of *S. aureus* metabolic activity (Figure 5G) confirmed a virtual elimination of biofilm on β-peptide-loaded coatings, compared to uncoated substrates and films lacking β-peptide. Overall, these results demonstrate that these β-peptide loaded coatings can prevent *S. aureus* biofilm formation on the surfaces of titanium substrates.

**Figure 5:**
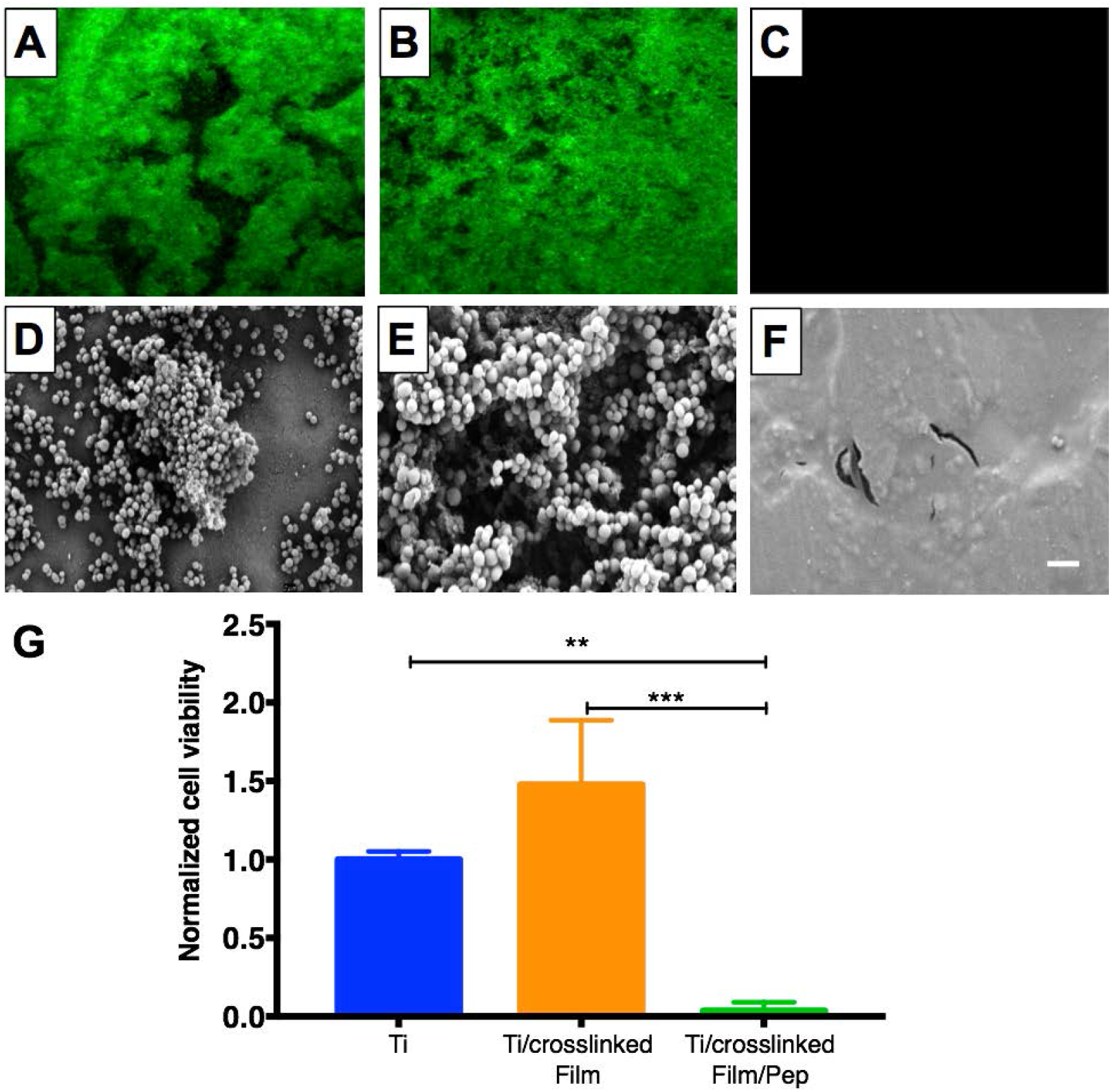
Evaluation *S. aureus* biofilm formation on titanium substrates coated with β-peptide-loaded CH/HA films. Uncoated, crosslinked CH/HA film-coated, and crosslinked CH/HA film-coated and β-peptide loaded titanium substrates were incubated for 24 hr with *S. aureus* cells (10^6^ CFU/mL) in biofilm inducing conditions (37°C, MH medium + 0.5% glucose). A-C) Representative fluorescence micrographs of GFP-expressing *S. aureus* strain AH1726 biofilms formed on the surfaces of A) uncoated, B) crosslinked film-coated (19.5 CH/HA bilayers) and C) crosslinked film-coated (19.5 CH/HA bilayers) β-peptide-loaded (0.44 mg/mL for 24 hr) titanium substrates. D-F) Representative SEM images showing the morphology of *S. aureus* biofilms formed on the surface of D) uncoated, E) crosslinked CH/HA film-coated and F) β-peptide-loaded crosslinked CH/HA film-coated titanium substrates. Scale bar: 10 µm. G) Quantification of live *S. aureus* cells on uncoated, crosslinked CH/HA film-coated, and β-peptide-loaded crosslinked CH/HA film-coated titanium substrates. *S. aureus* cell viability was quantified using an XTT metabolic activity assay and normalized to the uncoated control. Data points represent the mean values and error bars are the standard deviation of three independent experiments. Asterisks (*) indicate p < 0.01 by two-way ANOVA using Tukey’s multiple comparison test.

Following this proof-of-concept demonstration that β-peptide-containing coatings can prevent *S. aureus* biofilm formation in the short-term, we also evaluated their ability to resist *S. aureus* biofilm formation at longer time points, after some of the β-peptide had eluted from the films. For these experiments, we incubated titanium substrates coated with β-peptide-loaded films in PBS for up to 60 days, replacing the PBS solution every 2 days. At desired time points, substrates were challenged with an *S. aureus* inoculum and biofilm formation was assessed 24 hr later. Results shown in Figure 6A demonstrate that coatings loaded with β-peptide virtually eliminated the formation of *S. aureus* biofilms for up to 24 days after initiation of β-peptide elution. After 36 days, we also observed significant decreases in biofilm, with about 60% less biofilm on coatings loaded with β-peptide compared to bare titanium (Figure 6A). This extended inhibition of biofilm formation is consistent with the gradual release of β-peptides from the coatings over a period of 28 days (Figure 4).

Finally, we evaluated the ability of the β-peptide-loaded films to resist multiple bacterial challenges, as might occur after implantation of an orthopaedic device *in vivo*. We presented the substrates with five inocula of *S. aureus* cells for 24 hr each, washing the substrates between challenges (Figure 6B). As demonstrated in Figure 6C, the β-peptide-loaded coatings inhibited *S. aureus* biofilms after being challenged 5 times over a period of 18 days. However, the extent of biofilm inhibition was reduced for the last three challenges (~75% biofilm inhibition relative to uncoated control) compared to the first two challenges (~99% biofilm inhibition relative to uncoated control) (Figure 6C). Fluorescence micrographs of the biofilms formed on the surfaces of uncoated titanium surfaces, coatings lacking β-peptide, and coatings loaded with β-peptide confirmed the quantitative results reported in Figure 6C. Fluorescence micrographs acquired during the first two challenges demonstrate complete inhibition of *S. aureus* biofilms (Figure S6G and H) compared to uncoated substrates and control films lacking β-peptide (Figure S6A-B, D-E). However, following the fifth challenge we did not observe complete inhibition of biofilms on β-peptide-loaded coatings. In this instance, we observed the formation of less robust biofilms on β-peptide-loaded films (Figure S6I) as compared to uncoated titanium and control films lacking β-peptide (Figure S6C and F).

**Figure 6:**
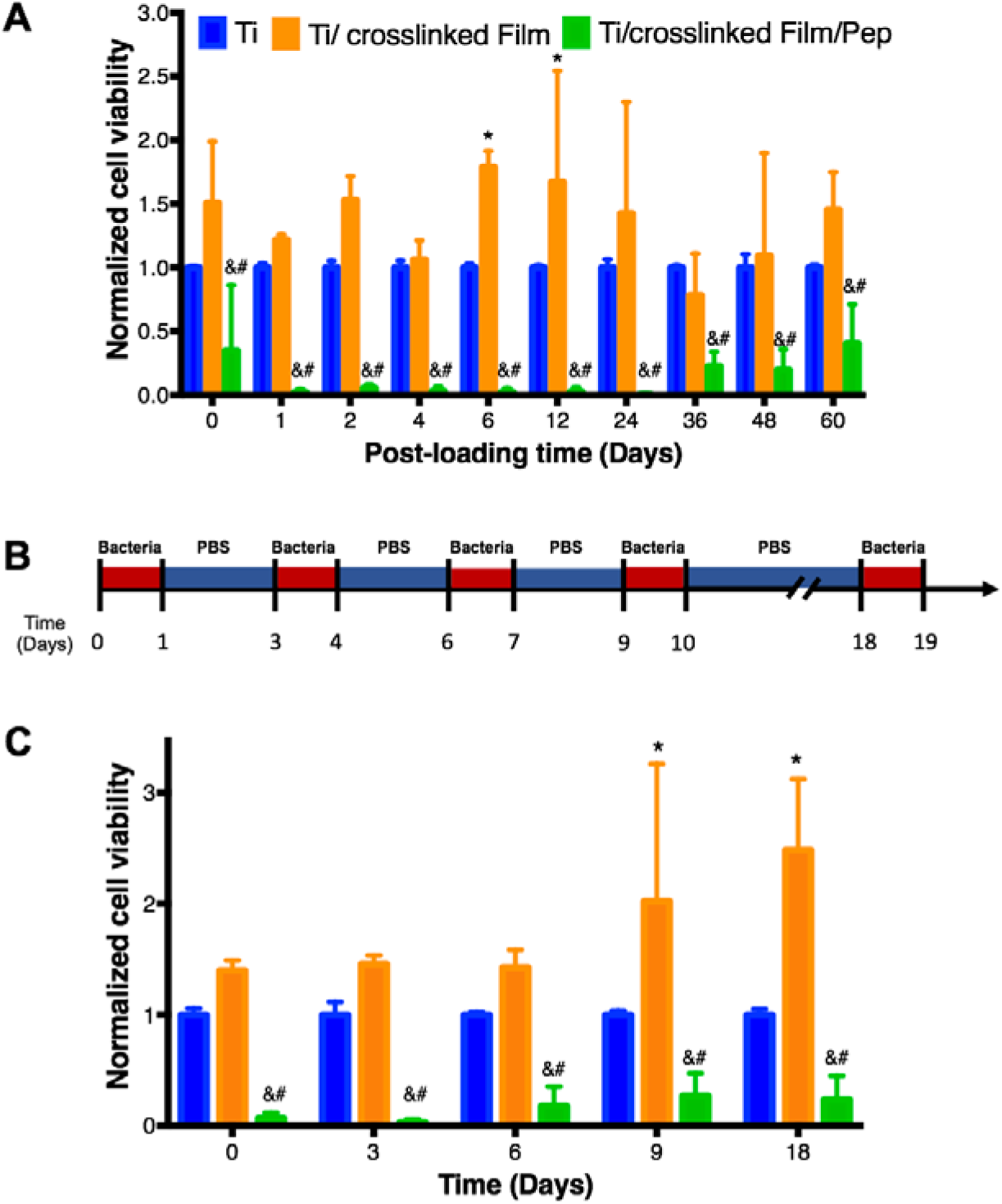
Quantification of *S. aureus* biofilm inhibition by β-peptide-loaded CH/HA films on titanium substrates after β-peptide elution in PBS for extended time and multiple short-term challenges. Uncoated, crosslinked film-coated (19.5 CH/HA bilayers), and crosslinked film-coated (19.5 CH/HA bilayers) and β-peptide loaded (0.44 mg/mL for 24 hr) titanium substrates were incubated for 24 hr with *S. aureus* cells (10^6^ CFU/mL) in biofilm inducing conditions (37°C, MH medium + 0.5% glucose). A) Long-term antimicrobial activity of uncoated, crosslinked film-coated and crosslinked film-coated β-peptide-loaded titanium substrates after being pre-incubated in PBS and challenged with *S. aureus*. B) Schematic for the protocol used when performing multiple *S. aureus* challenge experiments. Uncoated, crosslinked film-coated, and crosslinked film-coated and β-peptide loaded titanium substrates were challenged with an *S. aureus* inoculum for 24 hrs, followed by incubation in PBS for the specified period of time. These challenges were repeated until a total of 5 different challenges was performed. C) Antimicrobial activity of uncoated, film-coated, and β peptide post-loaded titanium substrates after multiple challenges with *S. aureus* inoculum. *S. aureus* cell viability was quantified using an XTT metabolic activity assay and normalized to the uncoated control. Data points represent the mean values and error bars are the standard deviation of three independent experiments. Asterisks (*) indicate p ≤ 0.05 between Ti and Ti/crosslinked film, # indicates p ≤ 0.05 between Ti and Ti/crosslinked film/Pep, and & indicates p < 0.01 between Ti/crosslinked film and Ti/crosslinked film/Pep by two-way ANOVA using Tukey’s multiple comparison test.

In summary, the results presented here suggest that crosslinked CH/HA films loaded with an antimicrobial β-peptide may be a promising approach for inhibiting *S. aureus* colonization and biofilm formation on titanium IFDs after implantation (Figure 5), with the ability to resist multiple *S. aureus* challenges and prevent biofilm formation for several weeks. These results may improve on the short-term antimicrobial activity of current coatings focused on titanium surface modifications for preventing *S. aureus* cell attachment [52,67,68] and demonstrate the effectiveness in preventing biofilms after eluting relevant antibiotic quantities in short time-periods (e.g., hours to days only) [22,52,54,69]. Additionally, our results demonstrate inhibition of *S. aureus* biofilms in a time frame (e.g., 3 months after surgical implantation) at which patients are most susceptible to microbial colonization. Therefore, our proposed therapeutic approach could potentially improve healing and further prevent implant failure in healthcare settings [15].

### 3.4. β-peptide-containing coatings elute β-peptide concentrations biocompatible with MC3T3-E1 cells

Many antimicrobial strategies have been reported for the prevention of implant-related bacterial infections. The adaptation of these strategies to orthopaedic implants should consider their biocompatibility with the bone microenvironment, including potential effects on bone cells [18,53,68]. Our results described above demonstrate that β-peptides in solution can prevent *S. aureus* biofilm formation with minimal toxicity against MC3T3-E1 cells. In addition, release of antimicrobial β-peptides from crosslinked films on titanium can prevent *S. aureus* biofilm formation. We next assessed the biocompatibility of our β-peptide-containing coatings incubated directly with MC3T3-E1 cells. The cytotoxicity of β-peptide-loaded films was evaluated by seeding MC3T3-E1 cells (5 x10^4^ cells/cm^2^) on the surface of uncoated titanium surfaces and PEM films loaded with β-peptide. MC3T3-E1 viability was quantified via a Cell Titer Glo assay after 24 hr. Our results demonstrate that films loaded with β-peptide (Figure 7, green bars) supported a similar extent of attachment of MC3T3-E1 cells after 24 hr compared to uncoated titanium (Figure 7, blue bars), suggesting that they are non-toxic to preosteoblast cells and exhibit good biocompatibility. Taken together with the results demonstrating *S. aureus* biofilm inhibition under these same conditions (Figure 5), these results indicate that β-peptide loaded films can be designed to elute β-peptide quantities that prevent *S. aureus* biofilms but do not cause substantial MC3T3-E1 cell toxicity.

We also quantified the viability of MC3T3-E1 preosteoblast cells on β-peptide-loaded coatings formed on titanium substrates after β-peptide elution in PBS for up to 60 days prior to MC3T3-E1 cell seeding. The viability of the MC3T3-E1 cells on films loaded with β-peptide was not significantly different than viability on bare titanium (Figure 7). When taken together, these long-term MC3T3-E1 viability results (Figure 7) and the long-term biofilm inhibition prevention assay (Figure 6A) demonstrate the selectivity of films loaded with β-peptide for inhibiting *S. aureus* biofilms without inducing MC3T3-E1 cell toxicity for prolonged periods.

**Figure 7.**
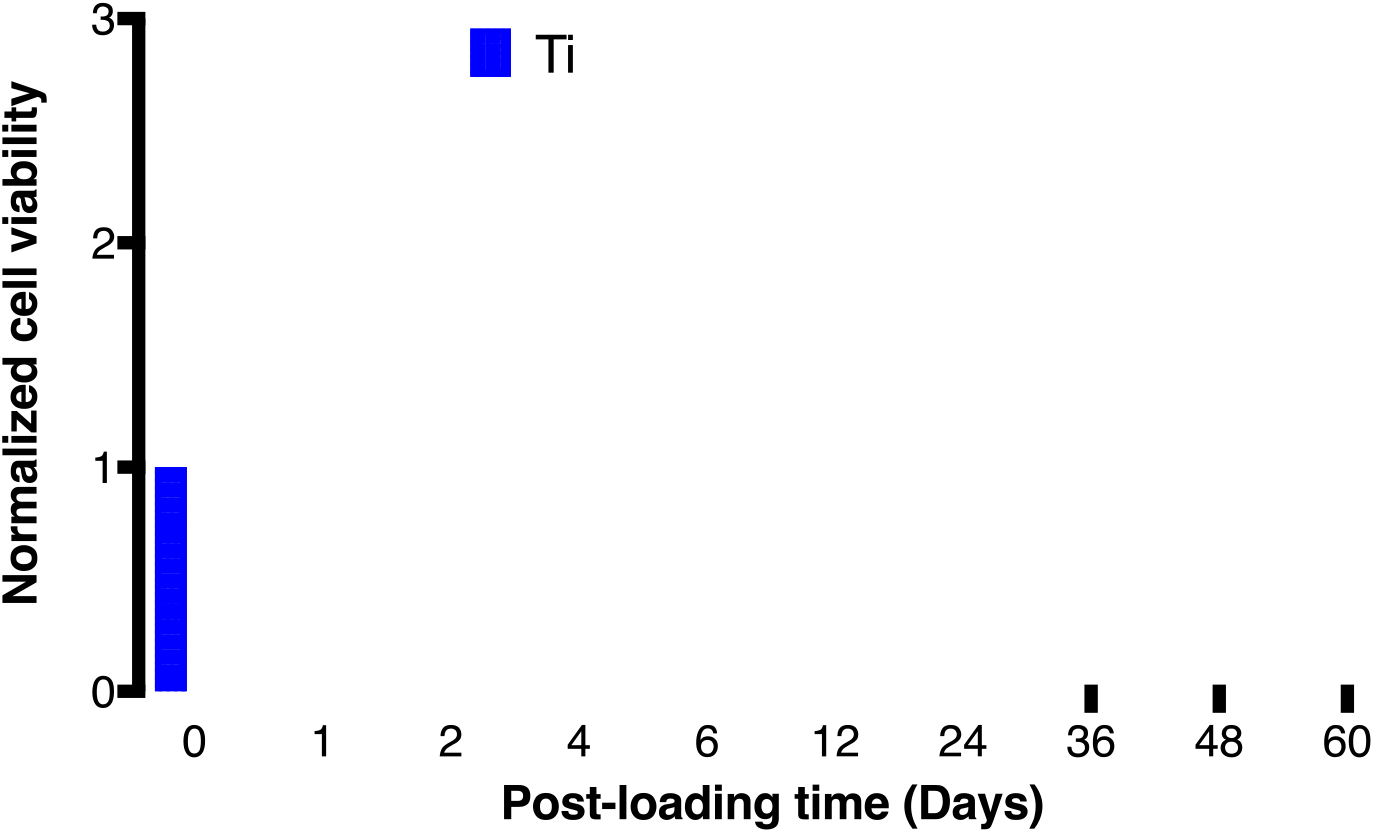
Evaluation of MC3T3-E1 cell viability on β-peptide-loaded CH/HA film-coated titanium substrates for extended periods of time. Uncoated, substrates and β-peptide-containing crosslinked CH/HA film-coated titanium substrates were incubated for 24 hr with MC3T3-E1 cells (5×10^4^ cells/cm^2^). Cell viability was quantified using a Cell Titer Glo assay and normalized to uncoated control. Data points represent the mean values and error bars are the standard deviation of three independent experiments. No significant differences were found between Ti and Ti/crosslinked film/Pep by two-way ANOVA using Tukey’s multiple comparison test.

## 4. Conclusions

This study used a layer-by-layer based approach to fabricate crosslinked CH/HA PEM films on titanium substrates. These films supported MC3T3-E1 preosteoblast cell attachment and proliferation for up to 6 days. We also demonstrated the incorporation of an antimicrobial β-peptide within the crosslinked CH/HA films to yield coatings that release β-peptide over a period of 28 days, which is relevant in the context of inhibiting bacterial attachment and biofilm formation over short to medium-term time periods. Furthermore, we showed that films loaded with β-peptide successfully prevented *S. aureus* biofilms formation *in vitro* without significantly decreasing MC3T3-E1 viability, suggesting promise for these films in the context of developing orthopaedic implant surfaces that resist biofilm formation. This result suggests promise toward developing novel strategies to inhibit biofilms without interfering with the osseointegration of IFDs. Moreover, β-peptide-loaded coatings inhibited *S. aureus* biofilm formation for up to 24 days and resisted five separate bacterial challenges over 18 days.

From this proof-of-concept demonstration, we conclude that crosslinked CH/HA PEM films loaded with an antimicrobial β-peptide are a novel and promising approach for inhibiting bacterial biofilms on IFDs and potentially reducing the occurrence of implant-associated infections in patients receiving IFDs. These promising results motivate further work to evaluate the ability of β-peptide-loaded films to inhibit *S. aureus* biofilm-related bone infections *in vivo*, as well as the development of more complex coatings (e.g., dual-delivery of antimicrobials and bone growth factors) that could also improve osseointegration. The coatings and strategies reported here also have the potential to be useful for inhibiting microbial colonization on the surfaces of other medical devices to potentially reduce the incidence of device-related infection in other contexts.

## 5. Acknowledgements

This work was supported by the National Institutes of Health grants 1R01 AI092225 and R21 AI127442 to S.P.P. and D.M.L. A. de L.R.L. was partially supported by an AOF research scholarship from the Graduate Engineering Research Scholars at UW-Madison. The authors gratefully acknowledge the use of facilities and instrumentation supported by the National Science Foundation through the University of Wisconsin Materials Research Science and Engineering Center (DMR-1121288). We thank Richard Noll for SEM training and help with FIB-SEM imaging. We also thank Benjamín J. Ortiz for his help with the PM-IRRAS film characterization and for many helpful discussions. Finally, we thank Prof. Alexander Horswill at University of Colorado for providing the GFP-expressing *S. aureus* strain (AH1756).

A. de L.R.L. and R.W. conducted experiments and collected data. M.R.L. synthesized and characterized the β-peptide. A. de L.R.L., D.M.L, and S.P.P. contributed to experimental design, data analysis and interpretation, and wrote the paper.

